# Uneven Walking Momentum Regulation Exhibit Different Strategies based on Age and State of Lookahead

**DOI:** 10.1101/2024.07.14.603456

**Authors:** Seyed-Saleh Hosseini-Yazdi

## Abstract

Uneven terrains enforce challenges to continuously regulate the step parameters. Since forward momentum contributes materially to the walking balance, its step-to-step regulation plays an important role to traverse terrain complexities. Here, we exhibited that young and older adults’ modulation were affected by the state of lookahead. With a normal lookahead, they demonstrated anticipatory control. The young adults encountered terrain irregularities similar to step-up mounting and dismounting in which the total mechanical work was minimum. On the other hand, the older adults might have put more emphasis on foot placement instead, as they slowed down before the encounters. Additionally, their control was extended beyond of young adults. On the other hand, with the restricted lookahead, the control was based on feedback. Since young and older adults’ momentum regulations were very similar, it also supported suggestions that older adults might inherently rely on feedback to control their gait.

## Introduction

Uneven walking imposes significant challenges due to diverse biomechanical and energetic factors that impact humans, especially older adults. It is well known that uneven walking requires excess energy expenditure compared to walking on even ground [1], [2]. Over uneven terrain, the walking stability is also poorer. Transitioning from even to uneven surfaces has been shown to decrease stability during locomotion [3]. The impact of uneven walking on momentum is also substantial. Since maintaining momentum is crucial for stability and efficient locomotion, particularly on challenging terrains [4], humans must adjust their walking patterns to effectively modulate momentum and prevent balance disturbances [5]. Therefore, more muscle activities are required [5]. The capacity to adjust momentum based on terrain variations is essential for overcoming the challenges presented by uneven surfaces and maintaining stability during locomotion [6]. Although it is demonstrated that young and older adults control their gait differently [7], nevertheless, there is limited information about their step-to-step center of mass (COM) velocity modulation over uneven terrains.

The uneven walking modulations are impacted by the state of lookahead. Matthis et al. [8] suggested that visual information is vital for the next foot landing selection, and humans need at least two steps or more of lookahead to approach even walking COM regulation. With the visual information about terrain perturbations, Darici & Kuo [9] have demonstrated that humans optimally anticipate and compensate for uneven steps by adjusting forward momentum and speed both before and after perturbation. This strategy involves speeding up before the perturbation and maintaining speed modulation over multiple steps post-perturbation. Additionally, the pre-activation reflexes can enhance a bipedal system’s ability to respond to unexpected perturbations on rough terrain [10].

The role of sensory inputs and the central nervous system in dynamically modifying control strategies to maintain gait stability and momentum in response to changing terrain conditions is important [3], [11]. Humans also recruit a common set of locomotor muscles at different speeds, for reactive responses to perturbations, and in anticipatory muscle activity before expected perturbations [12].

Without the visual information about the coming terrain when the lookahead is restricted, humans rely on feedback control [13]. It is suggested that the central nervous system flexibly modify control strategies to address unexpected changes in terrain morphology [3]. Therefore, the COM velocity (momentum) must be regulated instantaneously [14]. The exerted forces on the substrate and mechanical work performance therefore are essential based on feedback control [13]. Hence, if the momentum is controlled in fewer number of steps, the mechanical work performance elevates [13]. Prior experience with perturbations also enables humans make proactive adjustments based on estimation of surface conditions [15].

The young and older adults’ neuromuscular activation and gait modulation are different. During mid-stance at various walking speeds, older adults demonstrate greater activation of specific lower-extremity muscles, such as the tibialis anterior and soleus [16]. In older adults, increased coactivation across the ankle and knee joints are reported which perhaps is a compensatory mechanism to maintain stability during walking [16]. Alcock et al. [17] reported that older adults require greater activity from the primary motor cortex when walking at faster speeds. They proposed that this increased neural activity may reflect a higher demand for motor control and coordination to maintain gait stability. Additionally, it is noted that older adults generate smaller propulsive forces during push-of and for power generation, rely more on proximal leg muscles [18]. Since the proximal muscles are activated during stance phase to generate mid-flight works (after step-to-step transition) [19], [20], it might indicate that older adults are more reliant on feedback control. In contrast, young adults utilize a distribution of hip and knee extensor muscle force in early stance and ankle plantar flexor force in late stance to provide support and propulsion [21]. This indicates that young adults may rely more on feed-forward control mechanisms to generate propulsive forces without heavy reliance on feedback regulation. Moreover, Dingwell et al. [22] reported that while older adults may exhibit alterations in neuromuscular control and gait variability, young adults demonstrate more consistent stride-to-stride control. It also suggests a potential difference in the reliance on feedback mechanisms for gait modulation between the two age groups.

Walking speed modulation before and after encountering terrain perturbations is a complex process that involves various mechanisms to ensure stability and adaptability of human locomotion. Since in young and older adults, the state of musculoskeletal and central nervus systems are different, we expect they also demonstrate different traits in step-to-step COM velocity modulations. In this work we attempt to exhibit the potential influence of age and state of lookahead in controlling COM momentum over uneven terrains.

## Materials and Methods

We assessed the effect of age by hiring two groups of healthy adults to participate in our uneven walking experiment: Young Adults (age: 27.69 ± 5.21, mass: 71.0 ± 6.16 kg, six females and four males) and Older Adults (age: 66.1 ± 1.44, mass: 77.34 ± 12.13 kg, six males and four females). The procedure was approved by the University of Calgary Conjoint Health Research Ethics Board, and subjects provided written informed consent before the experiments.

To run uneven walking trials, we modified a split belt instrumented treadmill (Bertec Co., Columbus, Ohio, USA) to accommodate uneven terrains. The belt supporting structure was extended to provide the required room. We also fabricated uneven terrains by affixing construction foam blocks onto recreational treadmill belts [20], [23]. These foam blocks were arranged across the width of the belt, organized into sets measuring 0.3 m in length, with each set containing blocks of equal height. This configuration allowed either a flat foot landing or with a slight angle across two adjacent sets. The heights of the sets were 0 m (Even), 0.019 m, 0.032 m, and 0.045 m, respectively. Each terrain was identified according to its maximum foam block height (Max_h, Figure 1A). The terrains were open-ended and were wrapped around the instrumented treadmill’s original belt. We ran the trials at 0.8, 1.0, 1.2, and 1.4 m ⋅ s^−1^. The order of terrains and walking speeds were randomized. The trial schedule is provided in Table 1.

**Table 1:** The uneven experiment schedule: for Young (“YA” indicates normal lookahead and “ya” is for the restricted lookahead) and Older Adults (“OA”: is for normal lookahead and “oa” is for the restricted lookahead) for different terrains and speeds.

| Terrains Max_h(m) | 0 | Rough | 0.019 | 0.032 | 0.045 |
| --- | --- | --- | --- | --- | --- |
| Trials Velocities (m/s) |  |  |  |  |  |
| 0.8 | YA | YA | YA | ya-oa/YA-OA | YA |
| 1 | YA | YA | YA | ya-oa/YA-OA | YA |
| 1.2 | ya-oa/YA-OA | YA | ya-oa/YA-OA | ya-oa/YA-OA | ya-oa/YA-OA |
| 1.4 | YA | YA | YA | ya-oa/YA-OA | YA |

**Figure 1:**
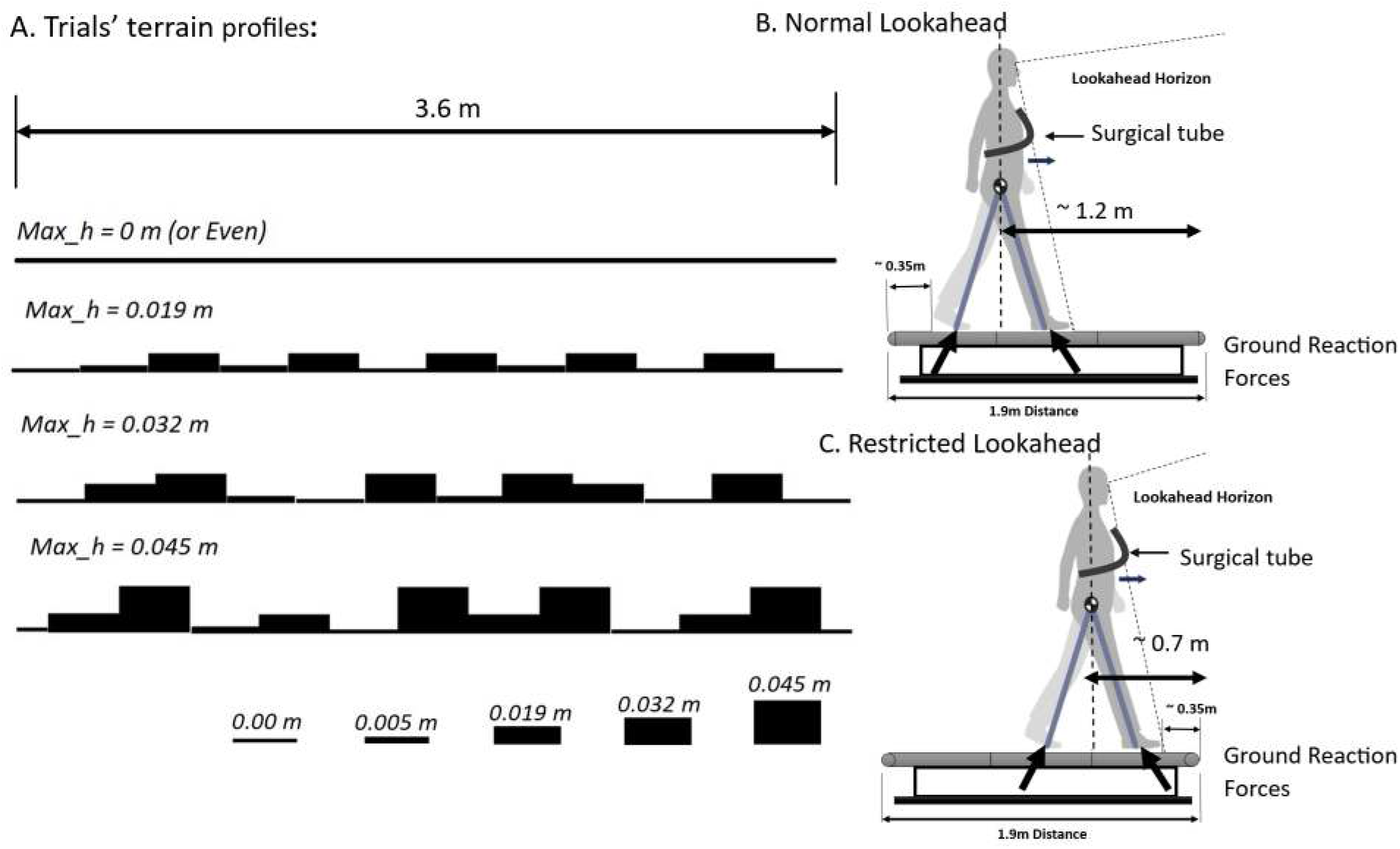
Experimental trials conditions: (A) The artificial uneven terrains’ profiles, (B) position for the normal lookahead in which subjects had almost two steps view horizon of the coming terrain, (C) position of the restricted lookahead where the subjects view of the coming terrain was limited.

To evaluate the impact of the lookahead, we implemented two conditions: *Normal Lookahead* (Figure 1B) in which subjects had almost two step length view of the coming terrain whereas in the *Restricted Lookahead* (Figure 1C), their view of the coming terrain was limited as much as possible. We help subjects remain in the designated positions with a surgical tube that was mounted across the treadmill with a slack. We asked participants to maintain gentle contact with the tube. We provided verbal feedback when it was necessary.

We collected ground reaction forces (GRF) at 960 Hz and low pass filtered them (cut-off 10 Hz). We derived the gait instances based on the GRF. We also calculated the COM velocity based on the GRF using the method outlined by the Donelan et al. [24]. For the step-to-step modulation we considered the COM velocities for mid-stances. The mid-stances were considered when the forward velocities were minimum for each stance.

We employed a standard marker set arrangement [20] and recorded motion capture using active markers (Phase Space, CA, USA, 960Hz). The duration of motion capture was the same as the GRF data recording. We calculated the step elevation changes as the difference between the consecutive averages of the minimum vertical positions of markers defining the first and fifth metatarsals (“*MTP*” metatarsal phalangeal joint).

We performed a regression analysis (Linear Mixed Effects Models, Statsmodels 0.15.0) to investigate the impact of step elevation changes on the COM velocity modulation when subjects encountered a certain perturbation (terrain amplitude change: COM Velocity = F(dE_i_). The dE_i_ represented step “i” elevation change concerning a certain perturbation encounter, i = 0: we considered the encounter to have happened at step “0”). The significant results were interpreted as indicative of step-to-step active modulations. Zero gains represented nominal magnitudes, while the positive gains indicated increases beyond nominal values. Likewise, the negative gains signaled declines below nominal magnitudes.

Since in the normal lookahead, subjects only had nearly two steps lookahead horizon, we considered the modulation to start after step -2. With the same token, we considered point of encounter as the start of regulation for walking with the restricted lookahead. Accordingly, we started our analysis from step -1 (i_start_ = −1) and step 0 (i_start_ = 0) for walking with normal and restricted lookahead, respectively. We ended our analysis at step +4 (i_end_ = +4, Figure 2).

**Figure 2:**
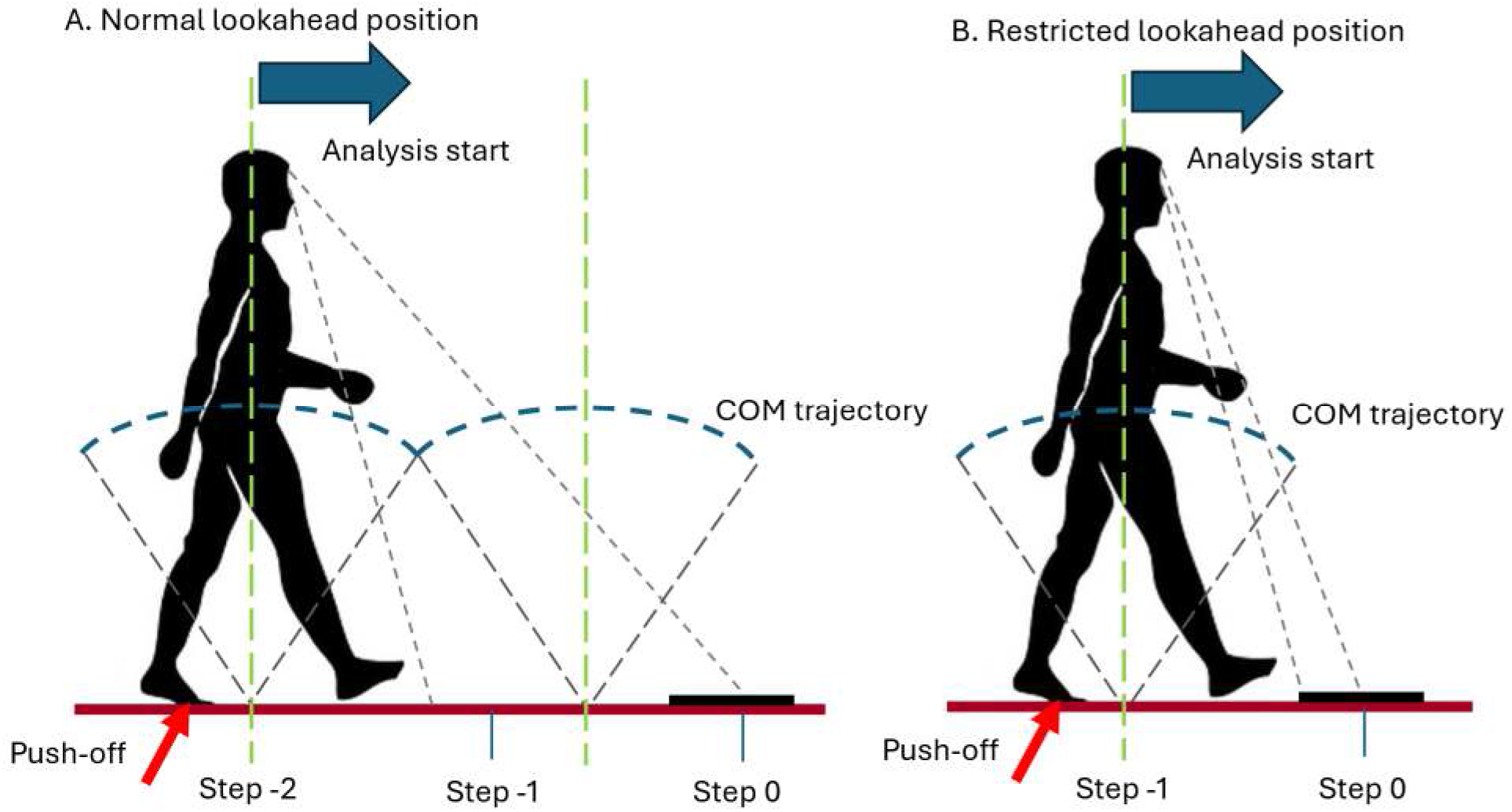
Step modulation based on visual information about the coming terrain (A) normal lookahead: since the sensory information was obtained at step -2, the analysis started from step -1, (B) restricted lookahead: since the sensory information was obtained at the point of encounter, the analysis started from step 0.

## Results

The conservation of momentum is crucial over challenging terrains to keep the balance. Therefore, we attempted to quantify the step-to-step COM velocity modulation during uneven walking by recording the step elevation changes and estimating the COM velocity based on the GRF. We also adjusted the lookahead horizon to examine our subjects’ strategies to negotiate the artificial uneven terrains.

The young adults with the normal lookahead depicted an anticipatory modulation [9] that started one step before the encounter by increasing their speed. At the point of encounter, their velocity declined which was associated to the conversion of kinetic energy to potential energy. Dismounting the perturbation, their velocity increased which was the inverse of going atop the perturbation. We assumed that the active modulation ended at step +2 since the COM reached the nominal magnitude again with no further changes.

The older adults also depicted anticipatory modulation that started at step -1. Nevertheless, it appeared that their modulation was different from young adults as they slowed down. Despite mounting the perturbation, their velocity increased slightly which might be associated to large positive mechanical works they must have performed going atop the perturbation [25]. Post encounter, the older adults’ velocity increased (step +1, and step +2). Since subjects walked on an instrumented treadmill with constant speed for each trial, the COM velocity decline in step +3 might have been to adjust their position to the assigned location. Likewise, we considered step +4 as the end of active modulation when the COM velocity reached the nominal value again. All in all, it is fair to say that older adults demonstrated a noisy velocity anticipatory modulation with the normal lookahead.

On the other hand, young and older adults depicted similar modulations with the restricted lookahead. Since the modulation started after the encounter, it must be based on feedback control trigger by other sensory information [26]. It seemed that since subjects walked on a treadmill with a constant velocity for each trial, they exhibited a tighter control than the normal lookahead, since they attain the nominal velocity at step +2. Therefore, their regulation was limited to only two steps (Figure 3).

**Figure 3:**
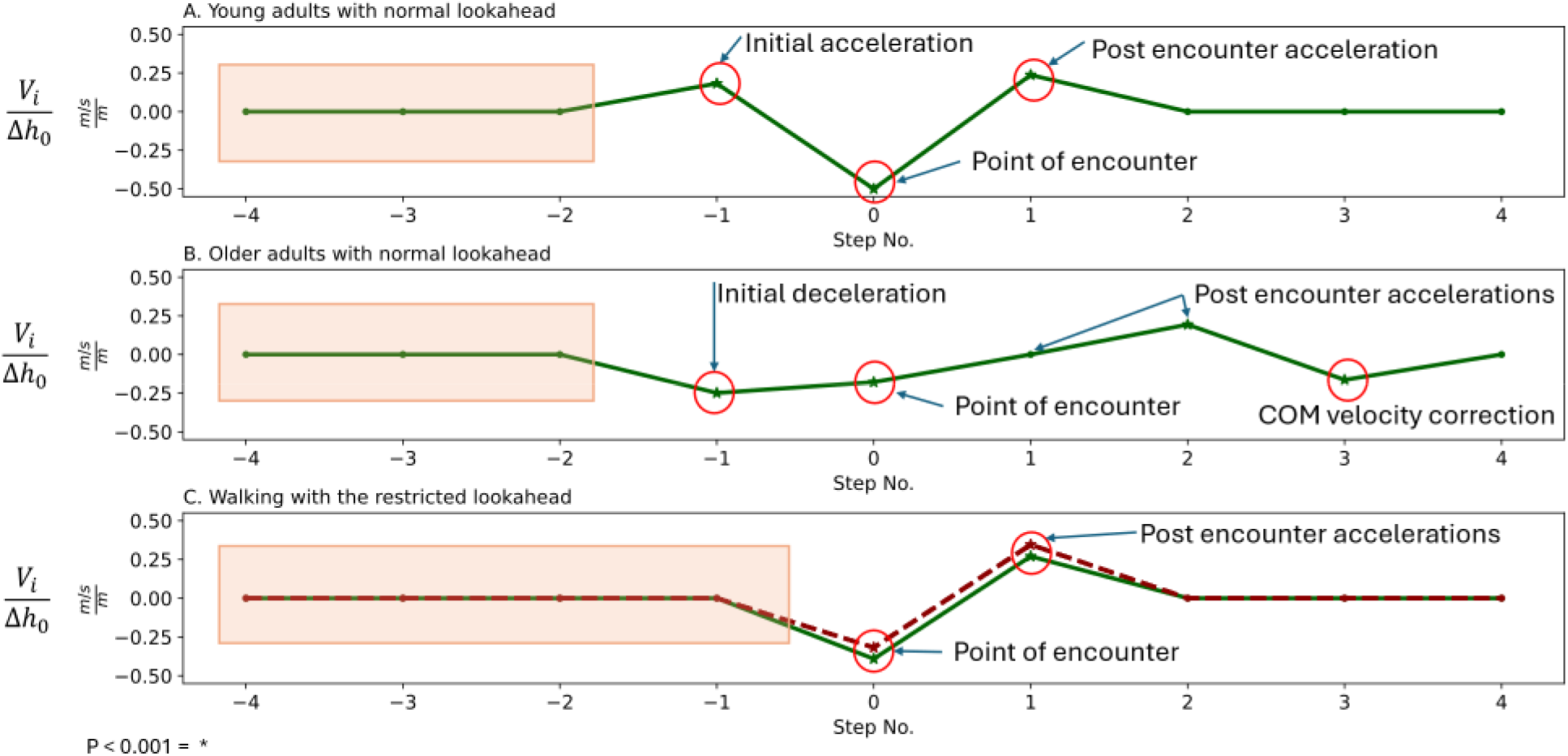
Step-to-step COM velocity modulation over the uneven terrains: (A) young adults with the normal lookahead, (B) older adults with the normal lookahead, and (C) walking with restricted lookahead: the solid line represents young adults’ regulation whereas the dashed line exhibits older adults. The significant modulations/regulations are represented by asterisks (*). The shaded areas were not considered for the analysis.

## Discussion

During uneven walking, the velocity modulation is crucial for adapting to varying terrains and maintaining stability in locomotion. It is proposed that visual information can induce changes in locomotion velocity and gait modulation [27], [28]. We have observed that to negotiate the challenges of the uneven terrain, young and older adults actively control and modulate their COM velocity. The state of lookahead appears to influence the way these two groups encounter perturbations. The COM speed modulation with normal lookahead suggests an anticipatory control while, with the restricted lookahead, the steep velocity reduction atop the perturbation and post encounter modulation indicate a control based on feedback.

With the normal lookahead, the young adults demonstrate anticipatory modulation when they receive visual information about the coming terrain [25], [29]. It is evident by the COM velocity increase before the encounter (step -1). Their speed profile suggests that uneven walking maybe decomposed into a series of perturbations mounting and dismounting [29]. On the other hand, the older adults’ anticipatory modulation started by a slowdown (step - 1). It might be a compensatory strategy to enhance stability and adaptability when facing environmental challenges similar to adopting wider steps over uneven terrains to maintain balance and prevent falls [30]. Older adults may prioritize attention on foot placement in response to perturbations stemming from impaired afferent feedback with aging [31]. It aligns with the observed slowing down before encountering terrain perturbations. The slowdown also might be a preparatory response to recruit additional motor units and adjust gait patterns to overcome perturbations effectively [32].

The deceleration before the perturbation encounter may pose some risks to older adults, however. Encountering terrain perturbations can lead to a loss of momentum, which poses significant risks for older adults by impacting their stability and increasing the likelihood of falls [4]. Falls are indeed a leading cause of injury among older adults, with most falls occurring during walking, particularly following perturbations of the support surface [9], [15]. The slowdown might necessitate a rapid repositioning of the base of support and suficient ground reaction forces from the musculoskeletal system to halt momentum [33].

Additionally, older adults exhibit greater force-generating errors and larger force variability when producing knee extensor eccentric control [10]. Since it is needed to exert large forces to go atop the perturbation [9], the slowdown exacerbates the magnitude of the positive work performance required for older adults. If they fail to generate the required positive work, it may lead to loss of balance and fall [4].

On the other hand, young and older adults demonstrate similar COM velocity controls with restricted lookahead. This suggests that older adults’ feedback control might be as efficient as that of young adults. Older adults often experience age-related changes in sensory processing and motor control [34], which can impact their ability to adapt to varying terrains efficiently. Age-related degenerative processes in the sensory and motor systems may lead to a shift from reliance on feedforward control to feedback strategies [35]. This shift could indicate that older adults may inherently rely more on feedback mechanisms to regulate their gait over uneven terrains, as feedforward strategies may become less effective in motor planning and execution [36].

It is also suggested that older adults, who may face challenges in feedforward control, could benefit from feedback mechanisms that provide real-time adjustments to maintain gait stability [37]. As such, instead of utilizing visual information about terrain perturbations, older adults might benefit more from environmental visual cues as feedback to maintain their momentum and stability [38].

In summary, we used our modified instrumented treadmill to study the gait and COM modulation over artificial uneven terrains with controlled amplitudes. One limitation of our experiment was the short treadmill length and the maximum height difference we could devise. As a result, we could detect a degree of correlation among the recorded step elevation changes (Figure 4). In perfect noisy terrain, only the gain for the point of encounter would be one while the rest are insignificant. Nonetheless, we could demonstrate the step-to-step COM velocity modulations based on the sensory information received in young and older adults. While young adults can utilize joint actuation that enables them to perform anticipatory modulations [21], the power generation redistribution in older adults [18] seems to be less efficient for feedforward control. Nevertheless, feedback control also requires visual cues from the environment other than the terrain perturbation [38], [39]. Since with anticipatory control, older adults exhibit slowdown before encountering perturbations, they might be prone to falling risk associated with the loss of momentum. Therefore, the suggested perturbation-based balance training that can enhance the reactive balance control of older adults, potentially reduces their fall risk. This improvement in reactive balance control aligns with the notion that older adults may excel in feedback mechanisms to adjust their gait in response to external perturbations, such as uneven terrains [40].

**Figure 4:**
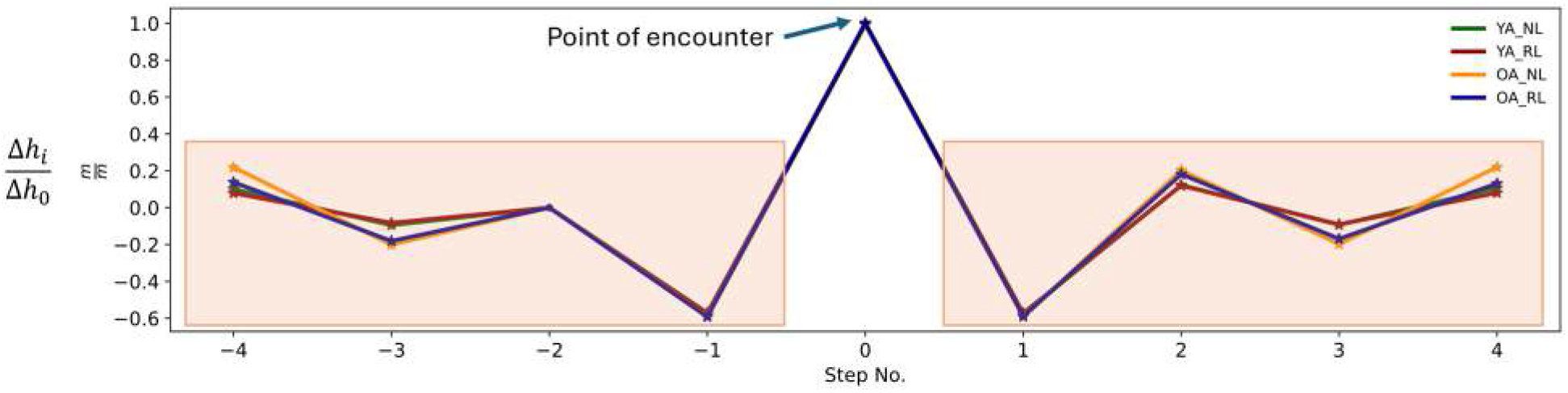
The step elevation for uneven walking: for the presented conditions, the step elevation gain at the point of encounter was 1 *m* ⋅ *m*^−1^. The significant step elevation changes with respect to the point of encounter (step 0) indicated a degree of correlation due to limited terrain length.

## Acknowledgements

This work was supported in part by the Natural Sciences and Engineering Research Council of Canada (NSERC) Discovery and Canada Research Chair (Tier 1) programs. The author would like to thank professor A.D Kuo.

